# Strategy for unlimited cycles of scarless oligonucleotide directed gene editing in *Chlamydomonas reinhardtii*

**DOI:** 10.1101/2024.01.18.576255

**Authors:** Ian L. Ross, Sabar Budiman, Hong Phuong Le, Dake Xiong, Fritz Hemker, Elizabeth A. Millen, Melanie Oey, Ben Hankamer

**Author notes:** All work reported here was carried out by the authors at the Institute for Molecular Biosciences, The University of Queensland, Brisbane, Australia. **Current email addresses and affiliations (in order of authorship):** IMB, The University of Queensland, Brisbane, Queensland 4072, Australia; IMB, The University of Queensland, Brisbane, Queensland 4072, Australia; Royal Botanic Gardens, Kew, Richmond, Surrey TW9 3AE, United Kingdom; Sichuan University of Science and Engineering, Zigong, China; Biochemie der Pflanzen, Heinrich-Heine-Universität, Düsseldorf; Ada Health Gmbh, Karl-Liebknecht-Straße 1. Berlin, Berlin 10178, Germany; IMB, The University of Queensland, Brisbane, Queensland 4072, Australia; IMB, The University of Queensland, Brisbane, Queensland 4072, Australia. **Conflict of Interest Statement:** The authors declare that they have no known competing interests.

## Abstract

CRISPR/Cas9 gene editing in the model green alga *Chlamydomonas reinhardtii* relies on the use of selective marker genes to enrich for non-selectable target mutations. This becomes challenging when many sequential modifications are required in a single cell line, as useful markers are limited. Here we demonstrate a cyclical selection process which only requires a single marker gene to identify an almost infinite sequential series of CRISPR-based target gene modifications. The *NIA1* (*Nit1, NR*; nitrate reductase) gene was this selectable marker. In the forward stage of the cycle, a stop codon was engineered into the *NIA1* gene at the CRISPR target location. Cells retaining the wild type *NIA1* gene were killed by chlorate, while *NIA1* knockout mutants survived. In the reverse phase of the cycle, the stop codon engineered into the *NIA1* gene during the forward phase was edited back to the wild type sequence. Using nitrate as the sole nitrogen source, here only the reverted wild type cells survived. By using CRISPR to specifically deactivate and reactivate the *NIA1* gene, a marker system was established that flipped back and forth between chlorate- and auxotrophic (nitrate) based selection. This provided a scarless cyclical marker system that enabled an indefinite series of CRISPR edits in other, non-selectable genes. Here, we demonstrate that this ‘*Sequential CRISPR via Recycling Endogenous Auxotrophic Markers* (SCREAM)’ technology enables an essentially limitless series of genetic modifications to be introduced to a single cell lineage of *C. reinhardtii* in a fast and efficient manner to complete complex genetic engineering.

## Introduction

Genetically editing specific targeted changes into the genome of living organisms is central to modern biology, and the discovery of Clustered Regularly Interspaced Short Palindromic Repeat (CRISPR) approaches greatly accelerated this. CRISPR employs ribonuclear protein (RNP) complexes consisting of short targeting guide RNAs and CRISPR-associated proteins (e.g. Cas9 (Charpentier and Doudna, 2013)). CRISPR/Cas9 mediated gene editing has many biological applications. These include algal biology, where biotechnological solutions are sought to advance renewable energy (Stephens et al., 2010), food and high value products and bio/nano materials (Karan et al., 2019; Ross et al., 2021). In some microalgal species CRISPR/Cas9 editing has been straightforward and successful, including *Nannochloropsis* (Wang et al., 2016) and *Phaeodactylum* (Nymark et al., 2016); in others, such as *Chlamydomonas,* it has been challenging, with low rates of transformation compared to other species, especially when transient expression of Cas9 and gRNAs were employed (Jiang et al., 2014; Guzman-Zapata et al., 2019).

*Chlamydomonas* is an important algal genetic model system, especially in photosynthesis and cilia biology (Harris, 2009). The quality of the *Chlamydomonas* genome sequence, the detailed understanding of its biology in comparison to other algal species, and the existence of large collections of well characterized mutants make it a preeminent model organism for genetic modification for biotechnological and renewable fuel-related applications, including the expression of recombinant proteins (Tran et al., 2013; Rasala and Mayfield, 2015; Scranton et al., 2015; Zedler et al., 2016; Ramos-Martinez et al., 2017; Baier et al., 2018).

Pioneering CRISPR/Cas9 editing work in *Chlamydomonas* by Shin et al. (Shin et al., 2016) and Baek et al. (Baek et al., 2016) employed transfected RNPs. Although the expression of Cas9 and guide RNAs from plasmids or genomic constructs has been achieved (Jiang et al., 2014; Greiner et al., 2017; Jiang and Weeks, 2017; Guzman-Zapata et al., 2019; Park et al., 2020), the RNP approach is simple to employ, more efficient, and has been widely used (Ferenczi et al., 2017; Greiner et al., 2017; Shamoto et al., 2018; Angstenberger et al., 2020; Cazzaniga et al., 2020; Dhokane et al., 2020; Kang et al., 2020; Kim et al., 2020; Picariello et al., 2020; Akella et al., 2021). Due to the relatively low CRISPR modification rates in *Chlamydomonas*, this approach typically employs marker systems, such as antibiotic selection genes, to enrich transformed populations prior to screening for gene edited cells. In *Chlamydomonas*, foreign transgenes such as selectable marker genes are typically inserted by non-homologous end joining (NHEJ) at the site of a double strand break, such as that produced at the target site by Cas9 (Baek et al., 2016; Shin et al., 2016). Importantly, Ferenczi et al. (Ferenczi et al., 2017) showed that single stranded oligonucleotides could be used for CRISPR-directed homologous recombination, with increased rates of successful modification. This has also been reported in other organisms (Hwang et al., 2013; Auer and Del Bene, 2014; Bialk et al., 2015) and the approach was recently used for targeted insertion of large inserts, up to 6kb (Kim et al., 2020).

The most successful and widely used selection markers in *Chlamydomonas* are exogenous antibiotic selection genes including *AAdA* (Cerutti et al., 1997), *AphVIII* (Sizova et al., 1996) and *Aph7* (Berthold et al., 2002) which, when transfected, confer resistance to spectinomycin/streptomycin, paromomycin and hygromycin respectively. Endogenous non-essential genes can also be used as *counter-selectable* markers, with mutants becoming resistant to toxic xenobiotics that kill wild-type cells. Examples include mutations in *APRT*(Schaff, 1994), *MAA7*(Palombella and Dutcher, 1998), *PPX1* and *ALS1* (Kovar et al., 2002; Akella et al., 2021).

Finally, auxotrophic strains including mutants in Arg7 (Purton and Rochaix, 1995), *Nic7*, *Thi10* (Ferris, 1995), *NIA1* (Kindle et al., 1989), *OEE1* (Mayfield and Kindle, 1990) and *SPD1* (Freudenberg et al., 2022) have been employed as selectable markers. As with antibiotic marker cassettes, transfection with a complementing gene usually involves random insertion creating a genetic “scar”. In a significant advance, Akella et al. (Akella et al., 2021) showed that by CRISPR targeting a selectable host gene along with the gene of interest, the use of co-transfected marker gene DNA could be avoided, leading to precise and scarless editing of the target gene.

Rapid, sequential editing to produce complex phenotypes is the next key challenge of CRISPR editing in *Chlamydomonas.* This would eliminate the need to use multiple markers or backcross single mutants to create multiply edited cell lines, for example to develop multiple simultaneous knockouts (e.g. to analyse complex traits), to introduce multiple point mutations (e.g. metabolic engineering) or to identify key amino acids in a target protein. This capability facilitates the creation of highly genome-modified algal cell lines to advance fundamental science and industrial production. Importantly, existing cell lines resulting from previous engineered genome changes could be rapidly modified to create new gene variants without the need to retrace engineering steps.

Here we report the SCREAM (*Sequential CRISPR via Recycling Endogenous Markers*) strategy; an efficient, convenient, reusable, scarless, fully reversible marker strategy which enables sequential addition of genetic modifications, without introducing fundamental changes to the original genotype. Using the same marker for all sequential target gene modifications establishes a standardized CRISPR technique that can be used to modify any set of target genes. We also sought a system that could be applied to a wide range of pre-existing strains and would not require the inclusion of a specific cassette or GMO construct into the cell line of interest.

Importantly, by regenerating the wild type selectable marker, SCREAM allows mutation of non-selectable target genes without the insertion of foreign marker DNA. The resulting organism is non-GMO in many global jurisdictions, with important commercial implications.

The SCREAM strategy requires a marker gene that enables both forward- and reverse-selection. It uses single-stranded oligodeoxynucleotide (ssODN) based editing to introduce (and then to reverse) a precise mutation (e.g., a stop codon) that flips the selection phenotype back and forth. This regenerates the original gene after each two-step cycle, and incorporates useful modifications that simplify screening.

The nitrate reductase (*NR; NIA1; Nit1*) gene was chosen as the selection system (Fernandez and Matagne, 1984; Fernandez et al., 1989). *NIA1* enables assimilation of nitrate as the sole source of nitrogen. Mutations in *NIA1* prevent *Chlamydomonas* from growing on nitrate, but *NIA1* mutants can be positively selected on media containing sodium or potassium chlorate (ClO_3_^-^) since the loss of NR activity confers resistance to chlorate (Prieto and Fernandez, 1993). Both forward (chlorate) and reverse (nitrate) selection strategies are consequently encoded by the one gene, *NIA1*. The single copy nitrate reductase gene *NIA1* has long been studied in the context of nitrate assimilation (Sanz-Luque et al., 2015) and was also the first gene used for nuclear transformation (and co-transformation) of *Chlamydomonas* (Fernandez and Matagne, 1984; Kindle, 1990; Blankenship and Kindle, 1992).

In forward SCREAM we employed an ssODN with Cas9 CRISPR RNPs to create a precise stop codon in the *NIA1* gene. Chlorate selection was used to enrich for *NIA1^-^* clones. For the reverse step (ssODN-mediated *NIA1* repair) we employed nitrate auxotrophic selection to identify genome edited clones in which repair of *NIA1* has occurred.

This approach enabled the use of a uniform selection assay strategy for any CRISPR-based target gene modification, provided a scarless marker and enables a limitless cyclical series of gene editing events. The widespread presence of nitrate reductase in other algal species and in land plants suggests that SCREAM using nitrate reductase offers a wide array of opportunities for plant biotechnology. However other endogenous dual selectable marker genes could also be employed.

## Results and Discussion

### Strategy: *NIA1* as an endogenous marker for SCREAM experiments

The *NIA1*-based gene editing strategy (Fig. 1) employs guide RNAs (gRNA) assembled with Cas9 to generate target specific ribonuclear protein (RNP) complexes. In the forward phase of SCREAM, guide RNAs are simultaneously directed at *NIA1* and the target gene(s) of interest. These are electroporated, with single stranded oligonucleotides (ssODNs) directed at *NIA1* (and, if desired, the target gene also) into *NIA1^+^ Chlamydomonas* cells. The “NIA1_Stop+BclI” ssODN contains mutations to introduce both a stop codon into *NIA1* (exon 2), and a new restriction enzyme site (*Bcl*I), to replace the natural restriction enzyme site (*Pfl*MI) in the original *NIA1* gene.

**Figure 1.**
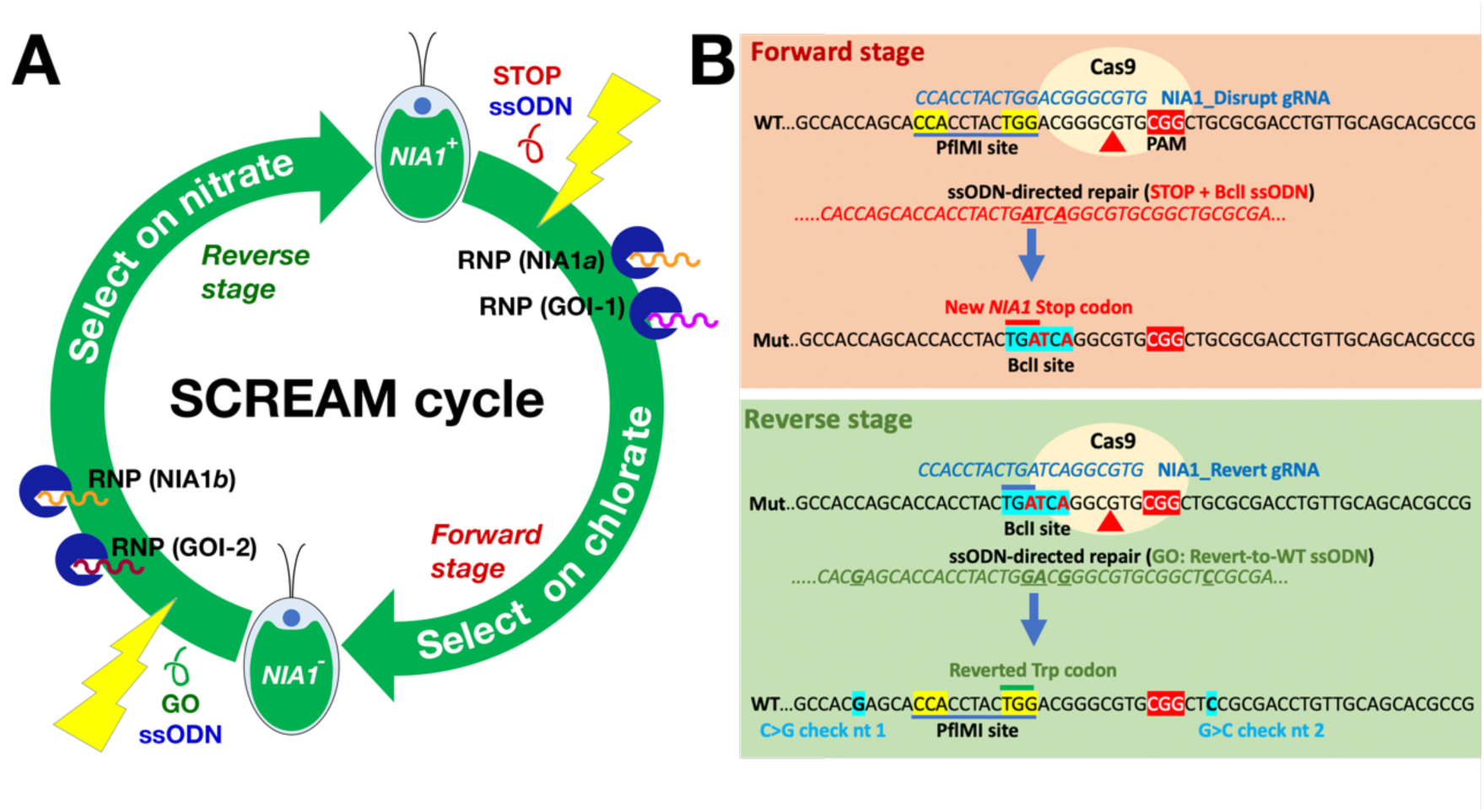
**A: SCREAM Strategy.** During the forward stage, *NIA1^+^* cells (top) are electroporated (yellow flash) with a STOP codon-containing ssODN as well as RNPs comprised of Cas9 (blue circles) bound to gRNAs directed against both the wild type *NIA1* gene (RNP NIA1a) and the first gene of interest (GOI-1). After selection for *NIA1^-^* cells on chlorate, cells containing both the *NIA1* stop codon (detected by BclI digestion of PCR amplicon) and the mutated GOI-1 (identified by gene specific PCR) are identified. In the reverse stage, the gRNA (NIA1b) targets the mutated *NIA1* and the GO (revert to WT) ssODN restores the wild type *NIA1* gene while the additional gRNA (GOI-2) targets the second gene of interest. The resultant edited *NIA1^+^* cells are able to grow on nitrate and can be selected on nitrate plates. Each cycle thus allows at least two genes of interest to be edited. **B: Detail of the ssODNs employed for homology-directed repair.** The *STOP+BclI* ssODN inserts a stop codon (TGG>TGA) that truncates *NIA1* translation in exon 2. It also eliminates a wild type *Pfl*MI site which relies on the TGG codon and installs a *Bcl*I site (TGA TCA). Nuclease digestion of PCR amplicons with *Bcl*I and/or *Pfl*MI thus enables rapid identification of *NIA1^-^* PCR amplicons likely to possess the ssODN sequence. In the reverse step, the *NIA1* Revert-to-WT ssODN reverts this altered site back to the wildtype sequence, restoring NR activity and nitrate-competent growth, ready to start the next round of alteration. The *Pfl*MI site is also restored and the *Bcl*I site is lost. For this work, two silent mutations 5’ and 3’ to the gRNA site allowed confirmation that the ssODN has been used as a template. These would not normally be required.

In the reverse SCREAM phase (i.e the “Revert to wild type” experiment, Fig. 1) the wild type *NIA1* sequence is restored, enabling growth on nitrate. The gRNA for reversion targets the introduced stop codon and *Bcl*I site, while the corresponding *NIA1* “revert-to-wt” ssODN eliminates the *Bcl*I site and re-establishes the wt gene sequence (including the wt *Pfl*MI restriction site). Additional gRNAs are simultaneously electroporated to mutate a second target gene(s) of interest. Consequently each SCREAM cycle generates at least two new sequential target gene mutations making sequential editing fast and efficient.

### *NIA1* deletion using CRISPR/Cas9

To validate the SCREAM process we first confirmed that our chosen gRNAs (Table 1) successfully directed *NIA1* gene editing (Appendix S1; Fig. S1) without the use of ssODNs. For this we employed the *NIA1^+^* CC-1883 strain (CC-1883) derived from the basic 21 gr wild type *NIA1^+^ Nit2^+^ Chlamydomonas reinhardtii* Sager strain (now designated CC-1690) which can grow on nitrate, with electroporation conditions initially calibrated using plasmid transformation (Fig. S2).

**Table 1.**
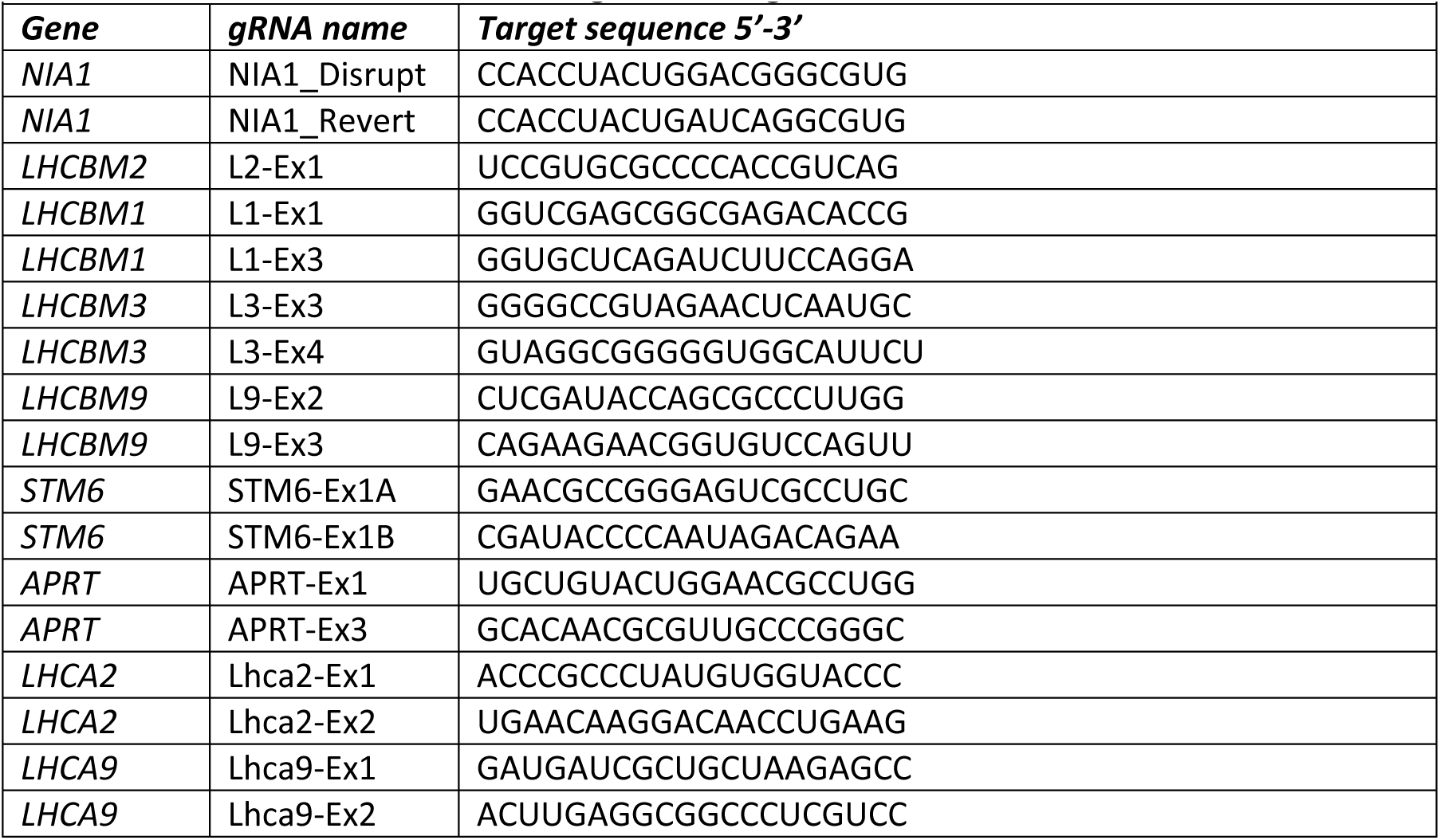
Guide RNAs used for CRISPR/Cas9 gene editing.

Following RNP electroporation and cell recovery, selection on chlorate resulted in 871 resistant colonies. After further selecting for clones unable to grow on nitrate (i.e. NIA^-^), PCR amplification of the *NIA1* locus, and *in vitro* Cas9 digestion of the amplicon (to identify loss of the gRNA site), 23 amplicons were chosen for sequencing. All 23 candidate mutants contained indels (Fig. S1d) which disrupted *NIA1* structure. Most (82%) were short deletions, in line with previous reports (Baek et al., 2016; Shin et al., 2016). Despite minimal sequence changes in some mutants, all grew poorly on nitrate suggesting that even small changes in the moco binding site (exon 2) disrupted *NIA1* function. Recently, Kim et al. (Kim et al., 2021) also showed that the *NIA1* gene was amenable to CRISPR modification in *Chlorella*.

### Specific ssODN directed *NIA1* mutation: Forward SCREAM stage

Having established that the chosen gRNA could successfully target *NIA1*, we next used ssODNs (Fig. 1B) to produce a specific targeted alteration to the *NIA1* gene. The ssODN incorporated a stop codon in exon 2 and introduced a *Bcl*I site for rapid identification of those PCR amplicons (Table 2) in which the ssODN was incorporated (Fig. 2). Meanwhile, at the protospacer adjacent motif (PAM) site a TGG→TGA stop codon replacement eliminated the native *Pfl*MI restriction site. Two additional base pairs were changed so that a *Bcl*I site was created (Fig. 1B). Consequently, in candidate mutants the *NIA1* PCR amplicon could be assayed by the presence of a *Bcl*I site, and/or loss of the *Pfl*MI site. Amplicon sequencing was therefore employed only for candidates which were likely to be correct, making the process fast and efficient. Most importantly, the completely defined nature of the mutation meant that it could be precisely targeted by a “reverse gRNA” and ssODN to revert the sequence to the wild type.

**Figure 2.**
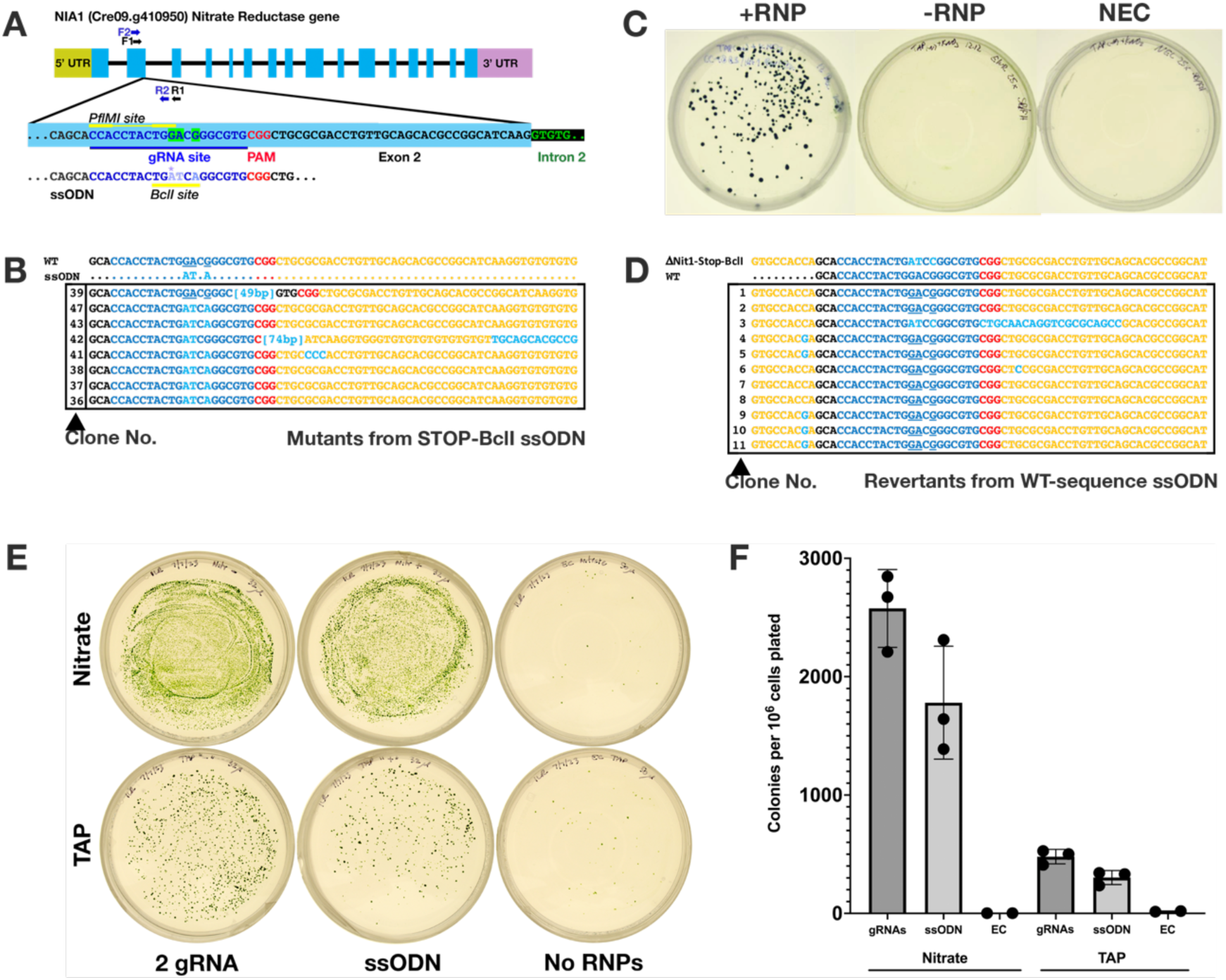
ssODN-mediated *NIA1* mutation and reversion. **A**. Diagram of the *NIA1* gene structure (top) and the expanded Exon 2 region sequence (bottom) including the gRNA target site (blue underlined), the PAM sequence (red) and the native PflM1 site (CCA *N5* TGG, yellow lines). PCR primers set 1 (F1, R1, black) and set 2 (F2, R2, blue) are indicated as arrows. Also shown is the ssODN (Bottom) used to introduce a specific mutation (*) designed to generate a stop codon and a *Bcl*I restriction site (TGATCA), while abolishing the native PflM1 site. **B.** SCREAM forward stage with ssODN: aligned PCR amplicon sequences of clones (No. 36-47) obtained after electroporation of CRISPR/Cas9 RNPs in a 50 μL reaction (10^8^ cells) coupled with the ssODN into the *NIA1* gRNA target region and plating on five chlorate selection plates. Note that six of the 7 mutants expected AT.A change. **C:** SCREAM reverse stage. Nitrate-utilizing (*NIA1*^+^) clones selected for on nitrate selection plates following electroporation of both a gRNA specific for the *NIA1*^-^ mutant sequence, as well as the wild type sequence ssODN which was used to revert the *NIA1*-Stop-Bcl1 clone to a wildtype *NIA1*^+^ phenotype. **D.** PCR amplicon sequences of clones obtained from ssODN-mediated reversion to the wild type phenotype. Note that 10/11 mutants (i.e. excluding Clone No. 3) have been edited back to the original wt GATCC sequence (light blue). **E.** Representative plates showing chlorate-resistant colonies resulting from growth in Nitrate Medium (NR induced) vs TAP (NR repressed), with either dual gRNA against the target gene or a ssODN and single gRNA. RNP plates represent one eighth of the electroporated cells. EC controls represent one quarter of the control cells. **F.** Colony quantification of dilution plates from Nitrate vs TAP experiment.

**Table 2.**
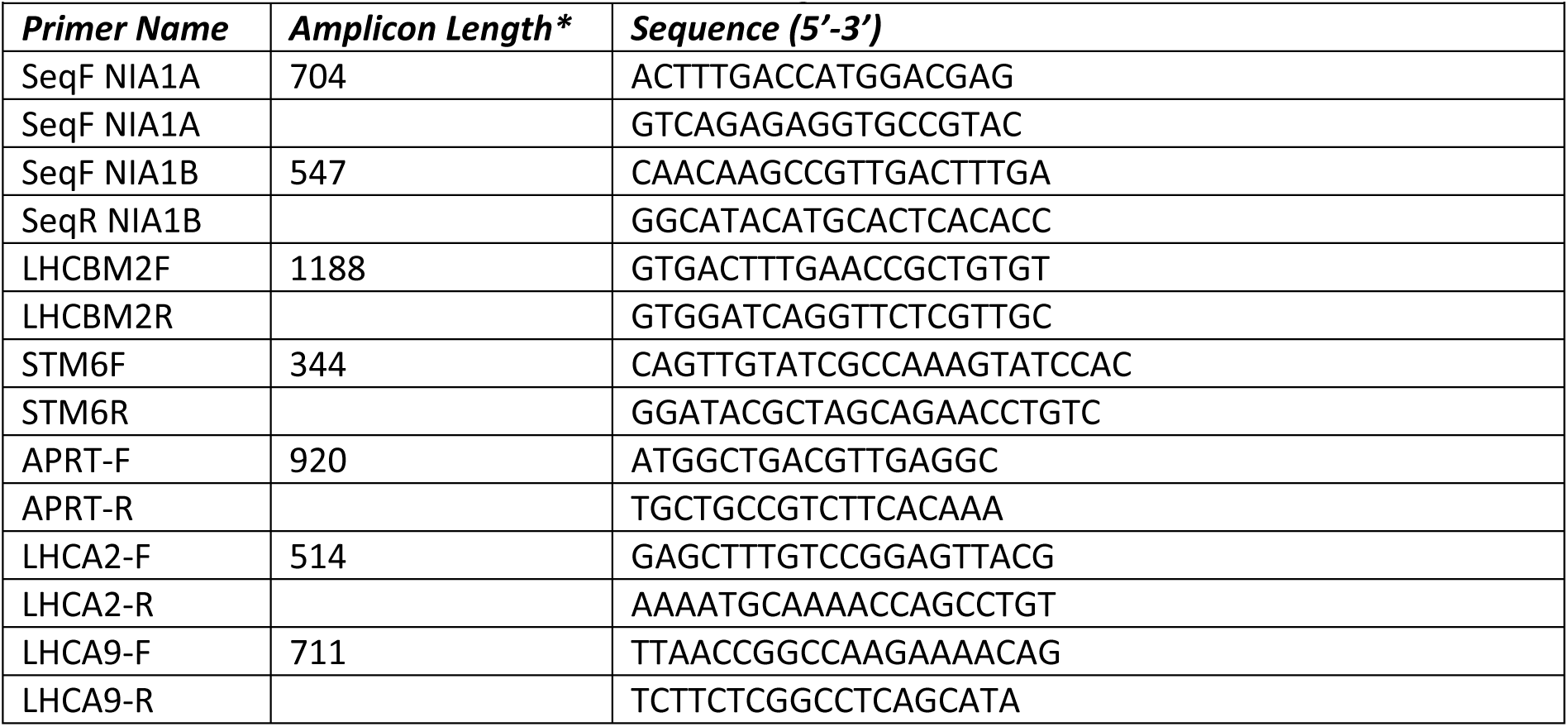
PCR primers used for amplification of edited regions.

The control (“no RNP”) plates showed ∼50 chlorate resistant colonies per plate (∼2.5 per million cells transfected). In contrast, colonies on plates transfected with RNPs and ssODN were too dense to be individually isolated. Therefore, an aliquot of cells was taken from each plate and diluted until single colonies could be identified. In this case, colonies from different plates likely represent independent clones. However, given the large number of colonies observed, even colonies from densely grown secondary plates are probably independent. All colonies (103 across the 5 plates) grew on TAP, but no colonies grew on nitrate, indicating that most colonies were CRISPR-induced, rather than spontaneous mutations (many of which can still grow on nitrate despite being chlorate-resistant; Appendix S2). To confirm this, genomic DNA was extracted, and eight mutants sequenced (Fig. 2A). One clone had a wild type *NIA1* sequence suggesting that it was a spontaneous mutant. All other clones contained the ssODN sequence at the gRNA target site (including the *Bcl*I restriction enzyme site) leading to premature truncation of nitrate reductase. Of these 7 ssODN modified clones, two had additional defects. One had a GCG>CCC mutation which would not prevent reversion via a wild type ssODN, while the other had an insertion difficult to revert to the wildtype sequence. Consequently 6/8 usable mutants were obtained for this forward SCREAM step. A clone with the desired ssODN correctly incorporated was chosen and designated the “*NIA1*-STOP-*Bcl*I” mutant for future experiments.

### Reversion of mutated sequence to wildtype: Reverse SCREAM stage

The *NIA1*-STOP-*Bcl*I clone chosen above was used to test the reverse SCREAM stage, that is, reversion to wildtype, via a WT ssODN (Fig. 3) and a specific mutant-targeted gRNA (Revert-to-WT gRNA, Table 3) targeted to the newly introduced target region of the chosen *NIA1*-STOP-*Bcl*I clone (Table 1). The ssODN included a silent G>C transversion on the 5’ arm (see Fig. 1) and a silent G>C on the 3’ arm, solely to demonstrate inclusion of the ssODN sequence.

**Figure 3:**
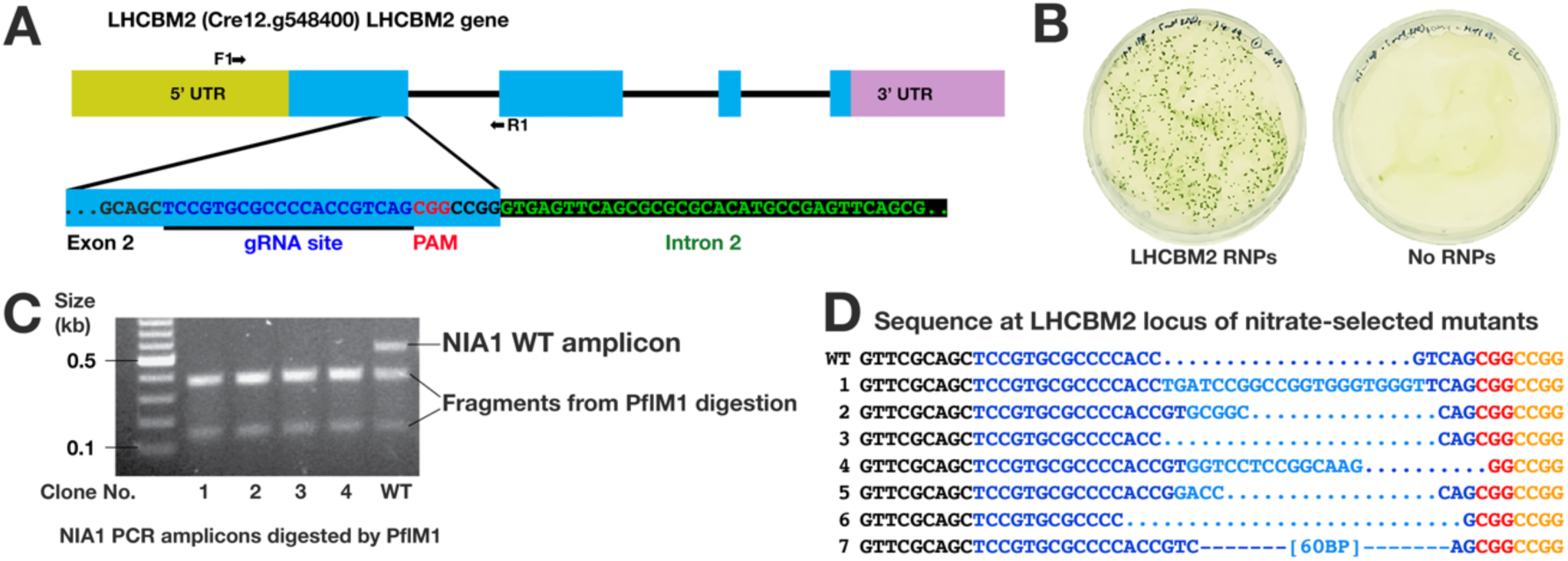
Use of SCREAM to target a non-selectable gene (*LHCBM2*). **A:** gRNAs targeting the *LHCBM2* gene. **B:** Transfection plates selected on nitrate for *NIA1* Reversion/*LHCBM2* knockout via SCREAM. Electroporation employed 100 µL of cell suspension (27.5 x10^6^ cells per reaction) and 50 µL RNPs, 5 nmol of LHCBM2 gRNA (Table 1). **C:** Clones tested for *Pfl*MI digested PCR amplicons. **D:** Sequence of wt and Δ*LHCBM2* knockout clones at the target site. Although not employed here, the 5’ transversion in the NIA1 gene also created a new restriction enzyme site (*BauI*) which offers an additional restriction enzyme based check for the presence of this sequence in the PCR amplicon.

**Table 3.**
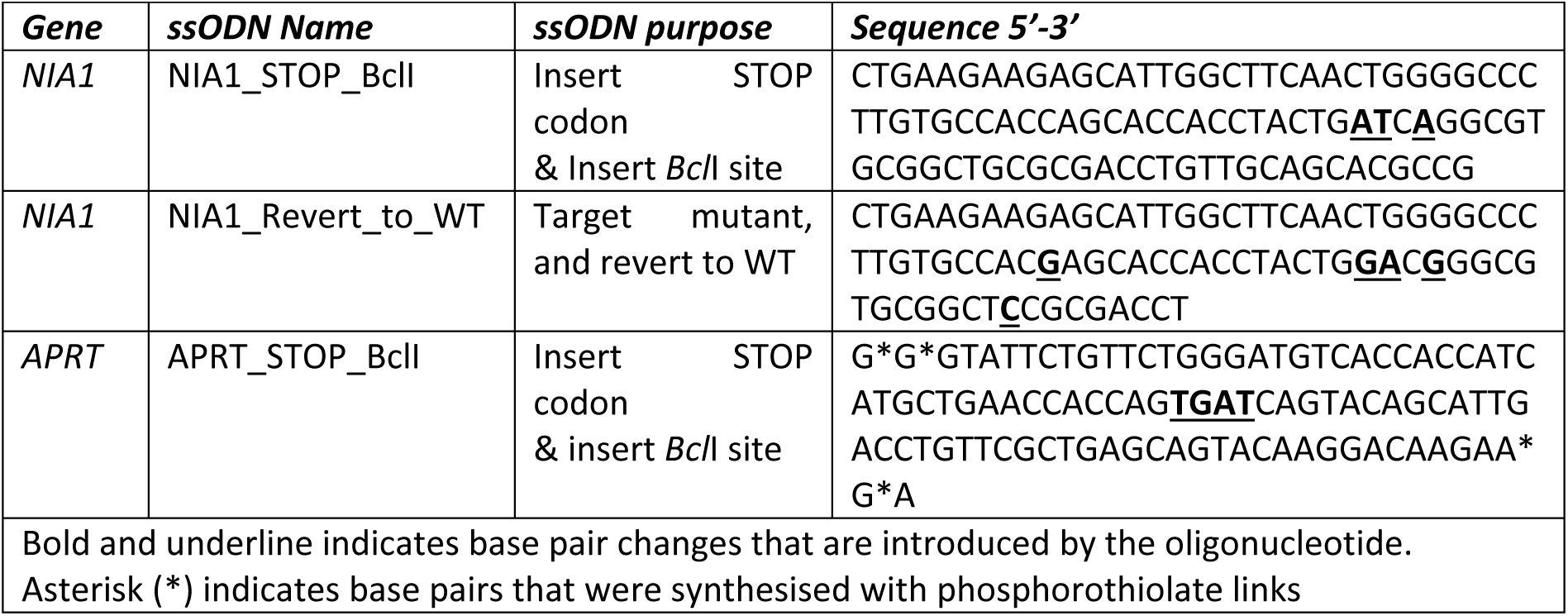
Single stranded oligonucleotides (ssODN) used to direct CRISPR/Cas9 gene editing.

The *NIA1*-STOP-*Bcl*I cells were grown in Nitrate Medium (TAP containing 5mM KNO_3_ instead of NH_4_Cl, and 2mM urea) as the nitrogen source for the *NIA1*^-^ mutants and electroporated in the same medium with 40 mM sucrose added. Although nitrate cannot be used by the *NIA1* mutants, it upregulates the *NIA1* locus via the transcriptional regulator *Nit2* (Fernandez and Matagne, 1986; Galvan et al., 1992), whereas ammonium ions (e.g. in TAP) would repress the *NIA1* locus (Sanz-Luque et al., 2013). An open locus is desirable for correct editing by Cas9 (Handelmann et al., 2023).

Following overnight incubation, the cells were replated on six Nitrate Medium plates containing only 5 mM KNO_3_ (no urea) to select for cells able to utilise nitrate (i.e. with an intact *NIA1* gene). As expected, no colonies occurred on the control plates, but the RNP plates contained many (Fig. 3B), suggesting that this selection protocol is very clean. Eleven colonies were checked by PCR followed by *Bcl*I digestion; all were *Bcl*I-resistant signifying alteration of the target locus from NIA^-^ to NIA^+^.

Sequencing revealed that 10/11 mutants contained the reverted NIA1 WT sequence (Fig. 2C), a success rate ∼90%. Clone 3 (Fig. 2C) retained the TGA stop codon and was out of frame. It is not known how this clone survived; potentially it has a restored *NIA1* gene but possesses a mutated PCR primer site, failing to amplify. Here, the observed amplicon may have been derived from genomic DNA from nearby non-surviving cells, or cells which temporarily survived on organic nitrogen from nearby dead cells. Of the 10 successfully reverted clones, 5 contained the upstream C>G silent base pair and one contained the 3’ G>C silent base pair change that confirmed the use of the ssODN as a template. The absence of clones containing both changes suggests that typically, ssODNs are partially digested during template guided repair. In summary, the forward (chlorate selection) step successfully resulted in ssODN incorporation and *NIA1* inactivation (∼60% success rate), whereas the subsequent reverse (nitrate selection) step successfully resulted in the genome sequence reverted to wildtype, with restoration of the *NIA1* gene (90% success rate).

### A complete SCREAM cycle with sequential modification of two target genes

Having established that our chosen gRNAs and ssODNs were able to drive a full SCREAM cycle with the *NIA1* gene, we next employed the cycle to co-select for CRISPR-induced mutations in specific *Chlamydomonas* genes for which mutant selection was otherwise difficult or impossible.

As part of another study into hydrogen production in algae, we required the production of a sequential modification of the genes encoding *LHCBM2* (*light harvesting protein B M2*) and *STM6* (also known as MOC1; *mTERF-like protein of Chlamydomonas-1*). The *LHCBM2* gene encodes the major light harvesting complex II protein 2 of *Chlamydomonas*, one of nine highly homologous major light harvesting genes of PSII (Sheng et al., 2019). Because of redundancy with eight other highly homologous LHCII genes, and an unknown phenotype, a screening assay for *LHCBM2* mutants was problematic. The *LHCBM2* mutant was also unavailable in the CLiP collection (Li et al., 2019) and was therefore created via CRISPR/Cas9 gene editing.

In contrast, the *STM6* (*MOC1*) gene knockout is the causative mutation in the *stm6* (*state transition mutant 6*) strain (Schonfeld et al., 2004). The *stm6* strain has a deficient mitochondrial alternative oxidase system (Wobbe and Nixon, 2013) and produces high levels of hydrogen (Kruse et al., 2005). We employed SCREAM to combine both mutations in a common CC-1883 background.

#### Target gene 1: LHCBM2

For the first target gene knockout, we transfected an anti-*LHCBM2* gRNA (Table 1; Fig. 4A) into the *NIA1*-disabled mutant created with the NIA1_Stop+BclI ssODN, so that the reversion-to-wildtype “reverse” SCREAM stage could be used for the first target gene modification. RNPs for the *LHCBM2* and *NIA1* target regions were assembled separately with Cas9, and then combined with the *NIA1* Revert-to-wt ssODN (5 nmol). These were transfected as before, and selected on nitrate-containing agar. No colonies were obtained when the RNP mix was omitted, whereas 3,380 colonies were obtained on the two RNP plates (Fig. 3). By comparison to TAP plates on which the electroporation mix was diluted (not shown), these 3,380 colonies represented around 0.56% of the entire population of viable cells. Twenty-two colonies were selected for PCR analysis. All amplicons were successfully cleaved by *Pfl*MI, suggesting restoration of the original *NIA1* wildtype sequence by ssODN-mediated repair. Sanger sequencing of the *LHCBM2* PCR amplicons of these clones revealed that 7 clones also contained edited *LHCBM2* target genes, yielding a target co-editing efficiency ∼32%.

**Figure 4.**
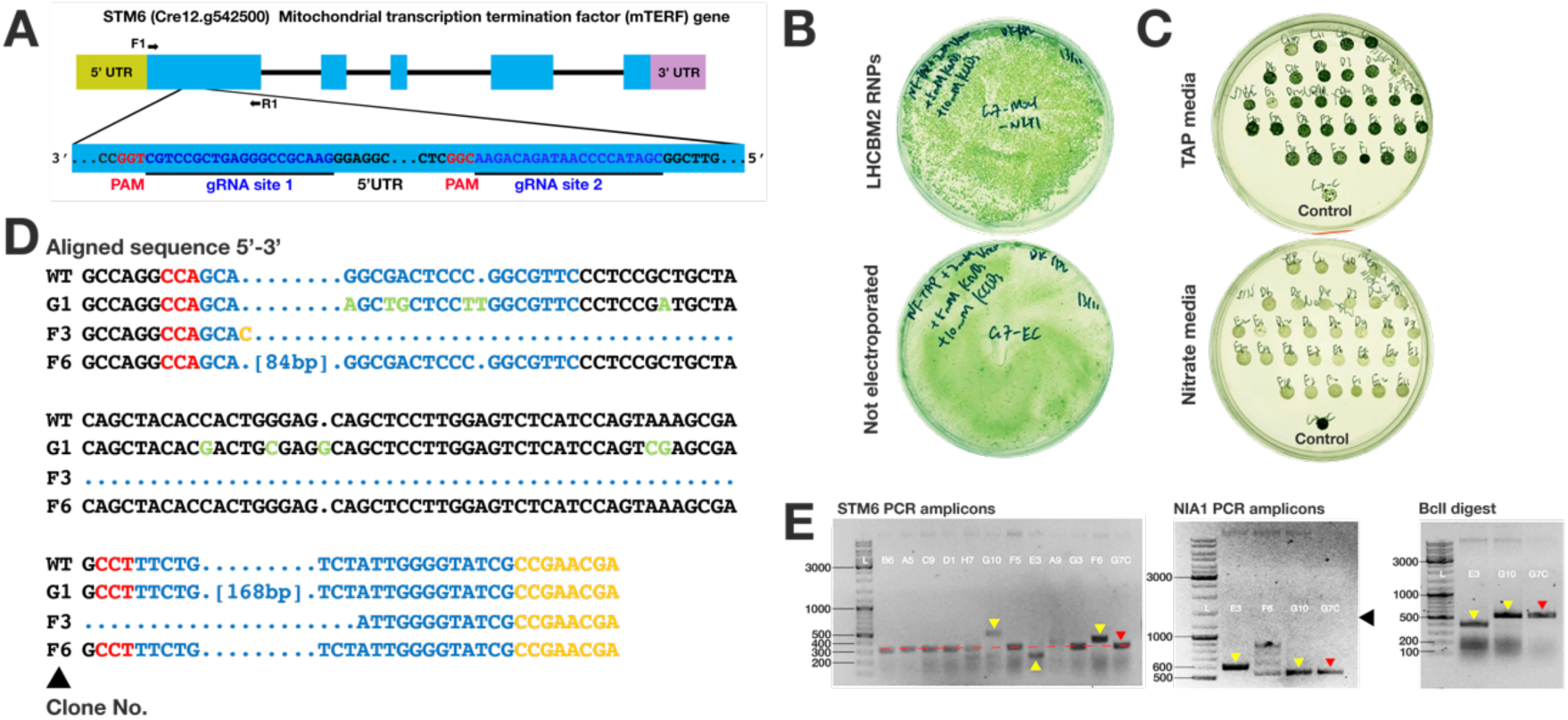
Use of SCREAM with dual gRNAs to sequentially target the *STM6* gene promoter in the *LHCBM2* mutant (*LHCBM2*/*STM6*). A: gRNAs targeting *STM6* exon 1. The use of two gRNAs for the STM6 exon 1 was designed to enable deletion of the chosen region just downstream of the start codon. Although Cas9 cleavage typically introduces indels, these are typically small in size and run similarly to the unmodified gene. In contrast, Cas9 cleavage at both sites produces a readily detectable change in the PCR amplicon size which is convenient for screening. **B:** Chlorate selection plates showing numerous candidate *NIA1*/*STM6* mutation colonies via SCREAM. **C:** Clones tested for growth on TAP vs nitrate plates showing loss of ability to grow on nitrate as a sole nitrogen source compared to control (wt) spot. **D:** Sequence of Δ*LHCBM2* Δ*STM6* clones. **E:** Analysis of clones by PCR; yellow triangles highlight altered *STM6* PCR amplicon size; Clone numbers in white; Red arrow indicates parent Δ*LHCBM2* clone G7.

Fig. 4 shows that the mutated *LHCBM2* genes had indels at 1 to 5 nucleotides from the 5’ end of the PAM site. In one case, part of the ssODN sequence used to revert the *NIA1* sequence to the WT, was inserted at the Cas9 cleavage site of the *LHCBM2* gene, illustrating a NHEJ-mediated knock-in effect. Of the mutants with successful indel formation at *LHCBM2*, all had the *NIA1* sequence returned to the wildtype sequence, with 5/7 containing the silent 5’ check nucleotide and 2/7 the 3’ check nucleotide, confirming that the supplied ssODN was used as the repair template. Although the *LHCBM2* KO was produced using a single gRNA, we routinely now employ two gRNAs per target gene for knockouts, since the resulting large change in PCR amplicon size is readily apparent, simplifying screening.

#### Target gene 2: STM6

In the previous step a CC-1883 *NIA1*^-^ mutant was engineered using SCREAM to simultaneously restore the *NIA1^+^* genotype and to knockout the *LHCBM2* gene. To further engineer a *STM6* (Δ*STM6*) knockout, a clone (G7) of the *LHCBM2* knockout cell line was then modified via forward SCREAM (*NIA1^+^◊NIA1^-^*), using chlorate selection, to create the triple mutant CC-1883 *NIA1^-^ LHCBM2^-^ STM6 ^-^* for a complete SCREAM cycle. Fig. 4A shows the *STM6* gRNAs employed. In this case we used a pair of gRNAs targeted against *STM6* exon 1, along with the *NIA1* gRNAs (Table 1) and the NIA1_Stop+BclI ssODN. The ability to immediately see a change in the amplicon size simplifies screening. As shown in Fig. 4B for one plate (of 3), many thousands of colonies resulted (estimated >5000 in total), far in excess of the chlorate spontaneous mutant background.

Four 96-well plates of colonies were picked (one from each of the four selection plates) into TAP medium, each containing 95 colony wells and one control (ΔLHCBM2 clone_G7) well. Colonies were transferred to agar plates containing nitrogen free TAP supplemented with 5 mM KNO_3_, to identify colonies disrupted in nitrate assimilation. As shown for one 96 well plate (Fig. 4C) no colonies grew effectively on nitrate, compared to the G7 (ΔLHCBM2 clone; see Fig. 3D) control. This suggests that the number of spontaneous chlorate mutants on the +RNP plates was very low and can be effectively disregarded. Around 300/396 colonies showed clear disparity between TAP and nitrate plates and were maintained, in case the mutation frequency of the target gene was low. Twelve randomly selected clones were checked by PCR amplification of the *STM6* target locus. Of these, 3 (i.e., ∼25%) showed large changes in the *STM6* PCR amplicon size, suggesting that either a large insertion occurred (amplicons larger) or that both gRNAs had caused double strand breaks in the genome (amplicons smaller), which had been repaired by NHEJ. Sanger sequencing of these three clones revealed that clones G10 and F6 contained insertions directly bridging the two gRNA sites, while E3 contained the desired deletion between the two gRNA sites. This supports the view that the altered PCR amplicon sizes were due to dual gRNA targeting events. Of the clones that did not show large changes in amplicon size, some will likely contain indels that also inactivate the target gene, although these were not sequenced. It should also be possible to create precise deletions by using bridging ssODNs.

These three clones were analysed for changes at the *NIA1* locus. Clone G10 contained a series of point mutations at the *NIA1* locus, E3 contained a large insertion, but F6 contained the correct STOP-*Bcl*I ssODN site and was therefore suitable for the next round of SCREAM gene modification.

### Reaction optimisation

As our standard CRISPR reaction generated far more *NIA1*-engineered clones than required to isolate a mutated target gene, the reaction was scaled down, to 5 million cells and one tenth as much RNP mix (see Methods). A voltage titration was conducted, estimating the actual yield of presumed CRISPR-edited *NIA1^-^*colonies, rather than employing plasmid-based transfection (Fig. S3). Electroporation of 25 µL of cell-RNP mixture in a 1 mm cuvette still yielded over 20,000 colonies (∼0.4% of cells electroporated) and was more economical. With electroporation, efficiency increases with voltage, but this is offset by increased cell killing at higher voltages (Fig. S3 B). The most efficient conditions represent a compromise between these two variables. Here, CC-1883 yielded most colonies at 200V 25 µF, infinite resistance, likely a reasonable setting for most *cw15* strains. However each strain (especially those with intact cell walls) will require electroporation calibration.

### *NIA1* locus accessibility is dependent on gene activation

Using chlorate-resistant colonies to estimate gene editing frequency, we compared the relative efficiency of NIA1 editing in CC-1883 cells grown in nitrate-containing, ammonium depleted media (in which the NIA1 locus is known to be induced) vs those grown in TAP, in which the presence of ammonium suppresses the NIA1 locus (Fig. 2E,F). For cells grown in nitrate media, ∼5 times as many colonies appeared compared to those grown in TAP, supporting the idea that gene activation (and thus chromatin accessibility of the locus) is an important factor in the frequency of gene targeting by CRISPR (for target genes as well as to the *NIA1* marker), suggesting that SCREAM should, if possible, be carried out under conditions where the target gene is actively being transcribed. Nevertheless, in targets examined to date, we have successfully obtained target gene modification even under conditions where the gene is expected to be repressed.

### Frequency of co-editing of target genes

An important variable in SCREAM is the frequency of target gene co-editing compared to the NIA1 marker, since editing frequency is dependent both on the target gRNA and the accessibility of the target site (Handelmann et al., 2023). The adenine phosphoribosyltransferase (*APRT*) locus was chosen as an independently selectable target so that frequency estimates could be performed simply using colony counting. *Chlamydomonas* cells with APRT knockout mutations can be selected on the toxic nucleoside analogue 2-fluoroadenine (2FA) since wildtype cells take up 2FA and incorporate it into their genomes, whereas mutants rely exclusively on *de novo* adenine synthesis (Littlefield, 1966). APRT had been originally considered as an alternative selection paradigm for SCREAM experiments but the selection of *APRT*^+^ cells requires treatment with both azaserine and alanosine to block the *de novo* pathway and simultaneous supplementation with adenine, which is a more complex system than the *NIA1* system. It is, however, a potential selective marker alternative to *NIA1* in the case of cell lines lacking several elements of the nitrate assimilation pathway. The *APRT* locus has already been successfully used in CRISPR experiments in *Chlamydomonas* (Guzman-Zapata et al., 2019) and *Chlorella* (Kim et al., 2021).

According to Ferenczi et al. (Ferenczi et al., 2017) inclusion of a single stranded oligonucleotide (ssODN) to direct homologous recombination increases the editing frequency of the target gene over use of a gRNA alone. We therefore compared RNP electroporations with and without *APRT*-directed ssODN, again using colony counting on selective media to score likely candidates, with the aim of improving the ratio of target (*APRT*) to *NIA1* mutants to improve screening efficiency.

Cells were plated on selective media comprising chlorate, 2-fluoroadenine or both, using a dilution series to generate plates with a suitable density for accurate colony counting, to estimate the mutational frequency of *NIA1* and *APRT* from the same electroporation reaction. Colony counts from control reactions lacking RNPs were used to measure spontaneous mutation rates for each condition but showed negligible numbers of colonies on single selection plates (<0.0005% of cells plated) and no colonies on double selection (2-fluoroadenine + chlorate) plates. The spontaneous mutant background can be therefore be disregarded.

On the most densely populated chlorate selection plates, ∼5x as many colonies occurred as on the corresponding double selection plates, suggesting ∼15-20% of cells were edited in both the *Nit1* and *APRT* loci (Fig. 5A,B). However, on the more sparsely spread double selection plates (where colony quantification is most accurate) the colony numbers, as a percentage of cells spread, were dramatically reduced. This suggests that single cells sparsely plated on selective media cope poorly with the simultaneous presence of both selection agents, even when genuinely edited at both loci. The true rate of double locus editing was thus presumed to be higher than indicated by the double selection plates. To improve quantification accuracy, 95 individual colonies were picked from single selection plates and gridded out on the opposite selection media, where the appearance of colonies identified clones that had both loci edited (Fig. 5C).

**Figure 5.**
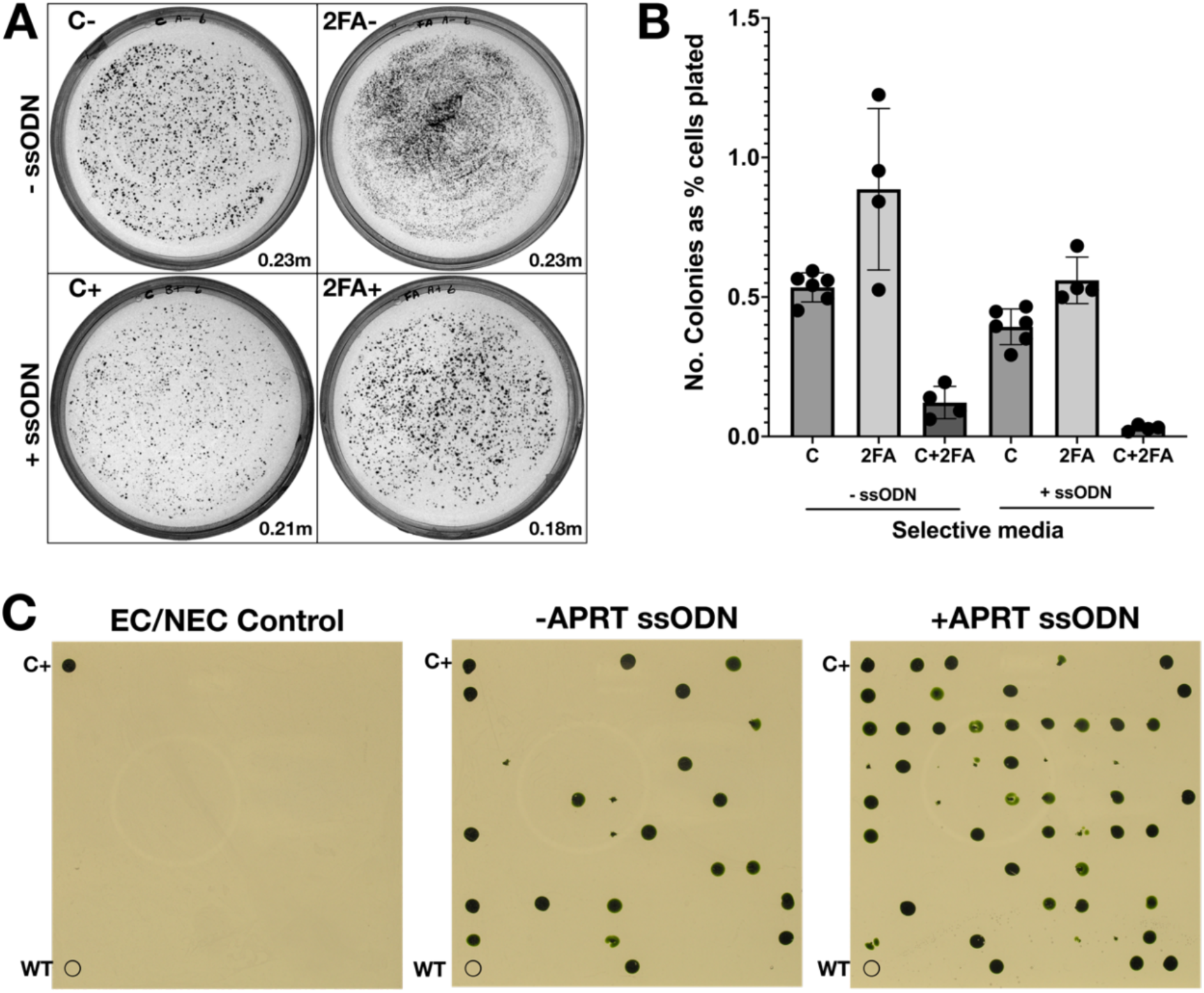
Use of ssODN to increase target gene modification efficiency. A: Representative selection plates showing RNP electroporation with (+) or without (-) *APRT*-directed ssODNs. Plates contained chlorate (“C”) or 2-fluoroadenine (“2FA”). Each plate represents ∼4% of each RNP electroporation (∼0.2 million cells; for specific values see bottom right-hand corner of each plate panel). Inclusion of the *APRT*-directed oligonucleotide led to a slight decrease in the total number of mutants obtained. **B:** Colony counting of the full dilution plate series was conducted to estimate the frequency of candidate *NIA1* edited clones (“C”: resistant to chlorate), candidate *APRT* edited clones (“2FA”: resistant to 2-fluoroadenine), and candidate *NIA1+APRT* doubly edited clones (“C+2FA”: resistant to both chlorate and 2-fluoroadenine). Clone numbers were estimated from selection plates using ImageJ. **C:** Randomly selected chlorate resistant colonies were picked, grown in TAP and gridded out on 2-fluoroadenine selective media. To validate the colonies on the grid, each plate contained a wild type CC-1883 negative control (“WT”) and a previously isolated *APRT* mutant positive control (“C+”). Randomly picked colonies consisted of chlorate-resistant colonies taken from plates spread with the non-RNP containing electroporated (“EC”) or non-electroporated (“NEC”) cells. (ii) the RNP electroporations without *APRT*-directed ssODN (“-*APRT* ssODN”) or (iii) RNP electroporations containing 0.5 nmol *APRT*-directed ssODN (“+*APRT* ssODN”).

None of the 95 chlorate resistant colonies isolated from control plates were resistant to 2-fluoroadenine (Fig. S4). In contrast, colonies derived from the CRISPR electroporation plates showed significant levels of cross-resistance (Fig. 5C and Fig. S4). From electroporation reactions lacking the *APRT*-directed ssODN, 16 out of 95 picked chlorate-resistant colonies were also strongly resistant to 2-fluoroadenine. When the *APRT*-directed ssODN was included in the electroporation, 39 out of 95 picked chlorate-resistant clones (40.6%) were simultaneously resistant to 2-fluoroadenine, demonstrating a 2.5-fold improvement in the efficiency of co-editing the *NIA1* and *APRT* loci. An independently conducted repeat experiment yielded a similar value of ∼41.6% (ssODN) vs 16% (2 x gRNAs) (Fig. S5).

This suggests that inclusion of the *APRT*-targeting ssODN led to a higher frequency of Δ*APRT* clones following chlorate selection, even though the absolute number of colonies obtained was somewhat lower (∼0.7x Fig. 2E) when the ssODN was included. Together with the higher primary colony numbers on single selection plates, this suggests that *APRT* is a more easily mutated locus than *NIA1*, possibly because the metabolic role of APRT ensures that the locus is usually open, whereas *NIA1* is an inducible gene and due to stochastic effects on gene expression, may be closed in a proportion of cells even under inducing conditions (Ross et al., 1994; Finn and Misteli, 2019). In the converse experiment, 2 out of 95 picked 2-fluoroadenine colonies were simultaneously resistant to chlorate when the *APRT* ssODN was not included, while inclusion of the ssODN led to 4 of the 95 2-fluoroadenine resistant colonies also displaying chlorate resistance. These results were lower than in previously range finding experiments (data not shown), potentially due to the variability of *NIA1* locus accessibility. In summary, these results suggest that using the SCREAM technique, electroporation of 5 million cells gives rise to around 25,000 *NIA1* candidate colonies, of which around 10,000 had the *APRT* target gene disrupted. Even allowing for undesirable disruptions (e.g., very large indels, badly placed indels at each locus, short indels that leave the reading frame intact) most of these will be effective knockouts, so that multiplexing (disruption of multiple target genes and/or library construction) is feasible as long as suitable (i.e. not PCR-based) screening for the target gene is available. In a single CRISPR ssODN-directed experiment, PCR screening of between 10 and 20 chlorate resistant colonies should suffice to identify a clone with a correctly edited *APRT* target gene. Broader screening will be required for targets with less accessible loci.

### SCREAM applicability

To date we have used SCREAM to produce single *Chlamydomonas* lines containing up to five consecutive gene modification events (first, KOs in NIA1, LHCBM1,2,3, and LHCA2; second, KOs in NIA1, LHCBM1,2,3, and LHCA9) and we have made at least fourteen engineered cell lines (listed in Table 4), which will be described in subsequent publications. We have trialled multiplexing SCREAM (i.e. the simultaneous use of gRNAs against more than one target). For example, strains 6 and 8 in Table 4 were produced within one electroporation. However, unless target gene conversion efficiencies are very high, the number of PCR reactions that need to be conducted to find the modified clones rapidly increases with the number of the genes targeted in a single mixed gRNA electroporation (see SI section D for further discussion). The simplicity of the SCREAM electroporation and plating process generally makes it easier to modify target genes in sequence, each requiring only a simple screening process, unless a library is specifically desired or a target gene-based screen (e.g. flow cytometry) is available. In contrast, multiple SCREAM experiments targeting a single gene each can easily be run in parallel.

**Table 4.** Summary of knock-out lines created using SCREAM to date.

| **Summary of knock-out lines created using SCREAM to date** |  |  |  |
| --- | --- | --- | --- |
|  | **Initial strain** | **Final genotype** | **KO method** |
| 1 | CC-1883 | *NIA1*<sup>-</sup> | **Indel** (*NIA1 Disrupt*) and separately using **ssODN** (gRNA *NIA1 Disrupt* + ssODN *NIA1_STOP_BclI*) |
| 2 | CC-1883 | *APRT*<sup>-</sup> | **Indel** (gRNAs *APRT-Ex1* + *APRT-Ex3*) and separately using **ssODN** (gRNA *APRT-Ex1* + ssODN *APRT_STOP_BclI*) |
| 3 | CC-1883<sup>1</sup> | *NIA1*<sup>-</sup>, *APRT*<sup>-</sup> | **Indel** (gRNAs *APRT-Ex1* + *APRT-Ex3*) and separately using **ssODN** (gRNA *APRT-Ex1* + ssODN *APRT_STOP_BclI*), with *NIA1* gRNAs and ssODN<sup>1</sup> |
| 4 | *NIA1*<sup>-</sup> | *NIA1*<sup>+</sup>, *LHCBM1*<sup>-</sup> | **Deletion** (gRNAs *L1-Ex1* and *L1-Ex3*) |
| 5 | *NIA1*<sup>-</sup> | *NIA1*<sup>+</sup>, *LHCBM2*<sup>-</sup> | **Indel** (gRNA *L2-Ex1*) |
| 6 | *NIA1*<sup>-</sup> | *NIA1*<sup>+</sup>, *LHCBM3*<sup>-</sup> | **Deletion** (gRNAs *L3-Ex3* and *L3-Ex4*) |
| 7 | *NIA1*<sup>+</sup> *LHCBM2*<sup>-</sup> | *NIA1*<sup>-</sup>, *LHCBM1*<sup>-</sup>, *LHCBM2*<sup>-</sup> | **Deletion** (gRNAs *L1-Ex1* and *L1-Ex3*) |
| 8 | *NIA1*<sup>-</sup> | *NIA1*<sup>+</sup>, *LHCBM1*<sup>-</sup>, *LHCBM3*<sup>-</sup> | **Multiplex deletion** (gRNAs *L1-Ex1* and *L1-Ex3*; *L3-Ex3* and *L3-Ex4*) |
| 9 | *NIA1*<sup>+</sup> *LHCBM2*<sup>-</sup> | *NIA1*<sup>-</sup>, *LHCBM2*<sup>-</sup>, *LHCBM3*<sup>-</sup> | **Deletion** (gRNAs *L3-Ex3* and *L3-Ex4*) |
| 10 | *NIA1<sup>-</sup>, LHCBM1<sup>-</sup>, LHCBM2<sup>-</sup>* | *NIA1<sup>+</sup>, LHCBM1<sup>-</sup>, LHCBM2<sup>-</sup>, LHCBM3<sup>-</sup>* | **Deletion** (gRNAs *L3-Ex3* and *L3-Ex4*) |
| 11 | *NIA1<sup>+</sup>, LHCBM1<sup>-</sup>, LHCBM2<sup>-</sup>, LHCBM3<sup>-</sup>* | *NIA1<sup>-</sup>, LHCBM1<sup>-</sup>, LHCBM2<sup>-</sup>, LHCBM3<sup>-</sup>, LHCA2<sup>-</sup>* | **Deletion** (gRNAs *Lhca2-Ex1* and *Lhca2-Ex2*) |
| 12 | *NIA1<sup>+</sup>, LHCBM1<sup>-</sup>, LHCBM2<sup>-</sup>, LHCBM3<sup>-</sup>* | *NIA1<sup>-</sup>, LHCBM1<sup>-</sup>, LHCBM2<sup>-</sup>, LHCBM3<sup>-</sup>, LHCA9<sup>-</sup>* | **Deletion** (gRNAs *Lhca9-Ex1* and *Lhca9-Ex2*) |
| 13 | CC-1883 | *NIA1<sup>-</sup>, LHCBM9<sup>-</sup>* | **Deletion** (gRNAs *L9-Ex2* and *L9-Ex3*) |
| 14 | *NIA1<sup>+</sup>, LHCBM2<sup>-</sup>* | *NIA1<sup>-</sup>, LHCBM2<sup>-</sup>, STM6<sup>-</sup>* | **Deletion** (gRNAs *STM6-Ex1A* and *STM6-Ex1B*) |
| <sup>1</sup>From Item 3 and onwards, *NIA1<sup>-</sup>* cell lines employ precise ssODN-mediate STOP codon insertion using the gRNA *NIA1_Disrupt* and the ssODN *NIA1_STOP_BclI*, while the *NIA1<sup>+</sup>* cell lines have a reversion to wildtype sequence which employ the gRNA *NIA1_Revert* and the ssODN *NIA1_Revert_to_WT*. The use of these *NIA1* gRNAs and ssODNs is implied but not stated explicitly. The gRNA names originate in Table 1.<br><sup>2</sup>These ssODNs have 2 phosphorothioate linkages at each end. |  |  |  |

Specific gRNAs against the target gene optimised using the many bioinformatic design tools available should generally suffice, especially if more than one is used. In the case of *NIA1* gRNAs, the relevant gRNAs for the forward and reverse stages are necessarily matched pairs, so optimal design for one gRNA may, in theory, result in less than optimal design of the other pair member. However, higher RNP activity against the *NIA1* locus may actually lead to a lower probability of <u>target gene</u> modification in the *NIA1*-edited chlorate resistant cells, so may not be desirable. In our view, at least as important as gRNA design is the need for the target gene locus to be accessible, for example by being transcriptionally active.

We expect nitrate reductase to be a suitable SCREAM marker for most microalgal species and we anticipate that the SCREAM approach will suffice for most routine CRISPR experiments to knock out or modify one or more endogenous genes, and to precisely insert transgenes at defined locations in the genome. All *NIA1^+^ Chlamydomonas* strains should be amenable, and some *NIA1^-^* strains (e.g. the *NIA1*-305 G-to-A point mutation (Plecenikova et al., 2013) can be readily repaired). Where *NIA1* repair is impossible, other dual-selectable markers genes such as *APRT* can be employed. While simplest to employ in haploid organisms it can also be used in diploids (e.g. *Arabidopsis*) especially where selective mutation or artificial generation of haploids (Shen et al., 2023) can reduce the endogenous marker to a single copy.

### Experimental procedures

#### Cell culture

*Chlamydomonas reinhardtii* strains CC-400 and CC-1883 were obtained from the Chlamydomonas Resource Center (St Paul, MN, USA) and maintained on 1% agar plates prepared with Tris-Acetate-Phosphate (TAP) medium (Gorman and Levine, 1965). Liquid cultures were cultured in Erlenmeyer flasks (∼120 rpm shaking) under white fluorescent light (50-100 µmoles photons m^-2^ s^-1^). Prior to electroporation, cells were grown for several days in Nitrate Medium, comprising TAP-N (i.e. TAP lacking ammonium chloride) supplemented with 5 mM potassium nitrate to induce the *NIA1* locus), with 2 mM urea to act as an alternative source of nitrogen for *NIA1^-^* cells.

#### CRISPR Guide RNAs (gRNA)

The nucleotide sequences of target genes were obtained from the Phytozome plant genome database using the *Chlamydomonas reinhardtii* genome version 4 (*NIA1*) and version 5.6 for *LHCBM2 (Cre12.g548400_4532)*, *STM6 (Cre12.g542500_4532)*, *APRT (Cre12.g548400_4532)*, and *SIRT6 (Cre12.g548400_4532)* genes (https://phytozome.jgi.doe.gov/pz/portal.html). Guide RNAs were designed using the Cas-Designer module of CRISPR RGEN Tools (Center for Genome Engineering, Institute for Basic Science, South Korea (http://www.rgenome.net/cas-designer/)). The parameters set in Cas-Designer for gRNA design were: up to 1000 nucleotides of a region of a target gene, SpCas9 from *Streptococcus pyogenes* (PAM and RGEN type) and with *C. reinhardtii v5.0* as the target genome. The gRNA was selected after consideration of the target position, out-of-frame score, and the number of mismatches. Cas-OFFinder (http://www.rgenome.net/cas-offinder/) was used to check the proposed gRNA for possible off-site targeting. In addition, design recommendations by Wang et al. (Wang et al., 2018) were taken into consideration. A list of gRNAs used in this work is provided in Table 1. Guide RNAs, tracrRNA and spCas9 enzyme were purchased from Integrated DNA Technologies as Alt-R® CRISPR-Cas9 crRNAs. Strain names and phenotypes are given in lowercase italics (e.g. *stm6*, *arg7, cw15*). Gene nomenclature follows the Phytozyme gene names, with commonly used former names provided in brackets upon first use e.g. *NIA1 (Nit1)*.

#### Single stranded DNA oligonucleotides (ssODNs)

The design of ssODNs for homologous template directed repair was based on the relevant gene at the site of the anticipated DNA double strand break created by *Streptococcus pyogenes* Cas9. All ssODNs for homology-directed repair were designed to include a ∼45bp “arm” on either site of the ds break site as recommended by Wang et al. (Wang et al., 2018). The ssODNs were unmodified DNA (*NIA1*) or had the end two base pairs modified with phosphorothiolate links for additional stability (*SIRT6*) and were purchased either from Integrated DNA Technologies or Sigma Aldrich (now Merck) and were rehydrated in nuclease free water to 100 μM and stored at -20°C. Sequences for the ssODNs employed here are given in Table 3.

#### Cas9 RNP In vivo mix preparation

For each guide RNA, an RNP reaction was prepared, and in the case of experiments with mixed sets of gRNAs these were combined after RNP preparation at the time of electroporation. To prepare the reaction, tracrRNA and gRNA (Integrated DNA Technologies; IDT) were first mixed in IDT duplex buffer (30 mM HEPES, pH 7.5; 100 mM potassium acetate) and heated at 95°C for five minutes. The solution was allowed to cool to room temperature to generate the RNA duplex. IDT duplex buffer and then Cas9 nuclease were added into the gRNA solution and incubated at room temperature for 15 minutes to allow RNP complex formation and then stored on ice. The final reaction contained, for each RNP used, 0.42 μM Cas9, 0.8 μM gRNA, 0.8 μM tracrRNA, and (if used) 20 μM ssODN. Where multiple gRNAs for a target gene were used, the total gRNA concentration was 0.8 μM. Cas9-gRNA RNPs for each gene were made separately and combined just before the electroporation stage.

### Electroporation

Electroporation was conducted using a Bio-Rad Gene Pulser XCell equipped with Capacitance Extender (CE) and Pulse Controller (PC) modules and employed 2mm or 1mm cuvettes.

Reaction mixes comprised 50 x 10^6^ cells in 200 µL Nitrate Medium with 40 mM sucrose, plus 50 µL RNP mix (or IDT Nuclease-free Duplex Buffer). Electroporation employed 2 mm electrode gap cuvettes (Bio-Rad) at 450 V, 25 µF, infinite resistance, exponential decay mode. The final scaled down protocol employed 5 x 10^6^ cells in 20 µL plus 5 µL RNP mix, electroporated with 1 mm gap electrodes at 200V, 25 μF, infinite resistance). After electroporation, cells were gently transferred to 10 mL Nitrate Medium for 48h in low light (15-25 µmoles photons m^2^ s^-1^) to allow gene editing to occur. For “non-electroporated controls” the cells were transferred to an electroporation cuvette, then out to recovery medium without a pulse being applied.

After electroporation, the cell suspension was maintained overnight in Nitrate Medium (including 2mM urea) to enable CRISPR editing to occur.

### Clone selection

After recovery, cells were pelleted at 600 x g, and resuspended in a defined volume of Nitrate Medium. Dilutions were made into sterile Nitrate Medium so that 140 µL was spread per plate. Selection plates (0.6% agar) were prepared one day prior to plating with a Nitrate Medium base medium. Positive selection was accomplished with the inclusion of a selection agent, either 7.5 or 10 mM sodium or potassium chlorate (Sigma Aldrich), or 8 µg mL^-1^ 2-fluoroadenine (FluoroChem). Negative (auxotrophic) selection was accomplished on plates containing 5 mM potassium nitrate but without urea. Colonies were picked with a sterile toothpick and placed in a suitable medium for expansion.

### Image processing and colony scoring

The plate image was separated into RGB channels using Adobe Photoshop (Adobe Systems Inc.) and a standard RGB setting was applied to enhance the contrast of the green colonies against the background (Red +185; Green -200, Blue +165) and images saved as monochrome images. For colony scoring, plate images were imported into ImageJ. The central region of the plate was isolated, and the outside (non-colony) part discarded. The contrast was set for maximum contrast between colonies and background and manually checked. A binary watershed filter was used to separate double colonies, and colonies larger than 16 pixels were quantified using the Analyse Particles function.

### Genomic DNA extraction

To yield high quality double stranded DNA, 5 mL of *C. reinhardtii* stationary phase-liquid culture was pelleted by centrifugation at 3000 × *g* for three minutes and resuspended in 500 µL DNA extraction buffer, pH 9.5 (0.1 M Tris-HCl, 1 M KCl, and 0.01 M Na_2_EDTA). The cell suspension was frozen and thawed five times, incubated at 75°C for 10 minutes, and cooled to room temperature. RNase A was added at a 10 µg mL^-1^ final concentration and the sample incubated at 37°C for one hour. DNA was then prepared by sequential extraction with 300 µL phenol:chloroform:isoamyl alcohol (PCI; Sigma Aldrich) then chloroform:isoamyl alcohol (CI; Sigma Aldrich), precipitated with 70% ethanol and resuspended in 100 µL of 10 mM Tris, 1 mM EDTA (TE) buffer pH 8.0. The genomic DNA concentration was measured using a Nanodrop 2000c (Thermo Fisher Scientific, USA) and stored in the -20°C-freezer.

### Chelex based genomic DNA extraction for PCR in 96 well plates

For single stranded DNA suitable for rapid PCR analysis, the method of Cao et al. (Cao et al., 2009) using Chelex resin was employed, including modifications described by Nouemssi et al. (Nouemssi et al., 2020) and Singh et al. (Singh et al., 2018).

### Agarose gel electrophoresis

DNA was analysed using 1 % (w/v) agarose gel electrophoresis at 75-100V in TAE buffer (40 mM Tris; 20 mM acetic acid; 1 mM EDTA) with 1x SyberSafe dye (Thermo Fisher Scientific, USA) included. GeneRuler DNA 100bp Ladder Mix (Thermo Fisher Scientific, USA) 3μL was typically employed as the marker, and DNA stained with SybrSafe dye (Thermo Fisher Scientific, USA) was visualised under UV light in a ChemiDoc XR system (Bio-Rad, USA).

### PCR Amplification

Genomic DNA extracted from CC-1883 (*cw15*, *nit+*, *mt-*) was used as a DNA template for PCR using primers with annealing temperatures listed in Table 2. The PCR product was examined using agarose gel electrophoresis or used as the template for Cas9 *in vitro* digestion, mutation genotyping by restriction enzyme analysis, or for Sanger sequencing.

### DNA sequencing

For Sanger sequencing, PCR products (3 µL) were incubated with their respective primers (10 µM) and sent to the Australian Genome Research Facility (AGRF, The University of Queensland, Brisbane, Australia). The resulting DNA sequence was analysed by pairwise and multiple sequence alignment using MUSCLE in SnapGene software (Dotmatics Limited) or by manual adjustment. Amino acid sequences were inferred and uploaded to Pfam (https://pfam.xfam.org/) for domain analysis. Amino acid sequences of mutated sequences were pairwise-aligned using BLASTP to identify altered amino acid residues.

#### Cas9 RNP *In vitro* Digestion

PCR amplicons from the *NIA1* gene of the first set of candidate knockout clones was evaluated for an intact Cas9 site by using Cas9-mediated digestion. The *in vitro* digestion was prepared for two control reactions, a wild type PCR amplicon combined with Cas9 only control (without gRNA), and gRNA only control (without Cas9). Amplicons from *NIA1* candidate knockout clones were then assessed using the RNP complex (1:2 ratio between Cas9 and gRNA). The ingredients for *in vitro* digestion were adapted from Ferenczi et al. (Ferenczi et al., 2017) and contained a pre-assembled RNP consisting of 600 nM Cas9 (IDT) and 1.2 nM gRNA (IDT), 1.2 µM tracrRNA (IDT), 1X NEB CutSmart Buffer (New England Biosciences), and 20 µL of PCR product in a final volume of 25 µL. PCR product and NEB CutSmart Buffer were added and IDT Nuclease-free Duplex Buffer used to adjust the final volume to 25 µL. All reactions were incubated at 37°C for 45 minutes, followed by heat inactivation at 65°C for 15 minutes. The fragments of the cleaved amplicons were separated and visualized using 1% agarose gel electrophoresis.

#### Restriction Enzyme screening of Putative Mutants

PCR amplicons were tested for the presence of specific restriction sites using restriction enzyme digestion. Restriction enzymes were purchased from New England Biolabs or Thermo Fisher and used according to the manufacturer’s instructions. Digests were visualized by agarose gel electrophoresis.

## Author Contributions

I.L.R., S.B., H.P.L., and D.X. secured funding and designed research; I.L.R., S.B., H.P.L., D.X., F.H., and E.M. performed research; M.O. contributed new reagents/analytic tools; I.L.R., S.B., H.P.L., and D.X. analyzed data; I.L.R. wrote the paper, and I.L.R., D.X., S.B., H.P.L, M.O. and B.H. critically edited/revised the manuscript.

## Acknowledgments

I.L.R. was supported by Australian Research Council Linkage grants LP170100717 and LP180100269. D.X. was supported by a University of Queensland PhD scholarship. H.P.L. was supported by an Australia Awards Endeavour Scholarship and by Can Tho University, Vietnam.

M.O. was supported by National Health and Medical Research Council Development Grant GNT1176216. We also thank Wiebke Weber for assistance with making the Lhcbm9 knockouts.

